# An experimental assessment of the distribution of environmental DNA along the water column

**DOI:** 10.1101/2020.11.30.402438

**Authors:** André O. Agostinis, Giorgi Dal Pont, Alexandre Borio, Aline Horodesky, Ana Paula da Silva Bertão, Otto Samuel Mäder Netto, Thiago Luis Zanin, Antonio Ostrensky, Marcio R. Pie

## Abstract

The study of environmental DNA (eDNA) is increasingly becoming a valuable tool to survey and monitor aquatic communities. However, there are important gaps in our understanding of the dynamics governing the distribution of eDNA under natural conditions. In this report we carry out controlled experiments to assess the extent and timing of eDNA distribution along the water column. A sample of known eDNA concentration was placed at the bottom of a 5-m high tube (20 cm in diameter and total volume of 160 L), and water samples were obtained at different depths over an 8 h-period. The presence of the target eDNA was assessed by qPCR analysis. This sampling protocol allowed for assessing the timescale for the diffusion of eDNA while minimizing the influence of turbulence. We demonstrate that, after a time-period of as little as 30 min, the eDNA had spread across the entire container. The implications of these results for eDNA sampling protocols in the field are discussed.

## Introduction

Environmental DNA (eDNA) is a method based on the detection of trace genetic material shed from organisms into their surroundings (Barnes and Turner 2016). Environmental DNA is composed by a range of particles, such as free DNA, organelles, cells, tissue fragments and metabolic waste (Turner et al. 2014; Wilcox et al. 2015). When suspended in an aquatic environment, this material can be sampled together with the water, extracted, and detected through molecular biology techniques (Ficetola et al. 2008). Surveillance and monitoring of aquatic species through eDNA is widely applied with advantages over traditional methods. This strategy, for example, can detect single or multiple species in one environmental sample (Harper et al. 2018), and the results can quantify relative biomass (Pilliod et al. 2014; Takahara et al. 2012). The main advantages of this approach are the shorter time requirements, increased cost-effectiveness, increased taxonomic resolution and non-invasive sampling (Eiler et al. 2018; Hunter et al. 2015; Thomsen et al. 2012). Several studies applied this method for aquatic organisms, such as fish (Miya et al. 2015), mussels and snails (Goldberg et al. 2013; Marshall and Stepien 2019), jellyfish (Minamoto et al. 2017), sharks (Bakker et al. 2017), amphibians (Pope et al. 2020) and arthropods (Toju and Baba 2018).

Although eDNA is a powerful technique, it is far from being standardized, as several methods are applied to capture and analyze field samples (Hinlo et al. 2017). Water samples, methods consist mostly of sampling the water column or the sediment (Buxton et al. 2017; Katano et al. 2017; Wittwer et al. 2018). However, studies rarely sample more than one depth, and when they do, the only parameter to compare them is detection rates or biodiversity (Andruszkiewicz et al. 2017; Yamamoto et al. 2016). Spread of eDNA horizontally was recently explored by studying flow from rivers (Jo et al. 2019; Pont et al. 2018; Sansom and Sassoubre 2017; Villacorta-Rath et al. 2020) but the vertical distribution is still fairly explored from a functional perspective. Vertical zoning is the structuring of communities through layers of species and communities across depths, which can potentially change dramatically in a matter of meters (Chappuis et al. 2014). eDNA concentration, composition and spatial distribution is then expected to vary as communities change through depth due to vertical zoning. While some studies conclude that there is a negligible impact on the detection and composition (Cordier et al. 2019; Currier et al. 2018; Eichmiller et al. 2014; Harper et al. 2020; Lafferty et al. 2020) others report differences (Andruszkiewicz et al. 2017; Cordier et al. 2019; Hänfling et al. 2016; Jeunen et al. 2020; Kuehne et al. 2020; Lacoursière-Roussel et al. 2018; Lor et al. 2020; Minamoto et al. 2017; Moyer et al. 2014; Murakami et al. 2019; Sigsgaard et al. 2020; Uthicke et al. 2018; Yamamoto et al. 2016; Zhang et al. 2020). These studies, however, vary significantly between the water body architecture, water composition, depth sampled, sampling strategy, target organism, detecting technique, extraction protocol and molecular marker. This leads to an inconsistent pattern that can be interpreted as detection relies more on the organism’s biology than depth (Minamoto et al. 2017). However, these studies do not consider how eDNA moves through the column as both (a) the sources of biological material are still in the water body, releasing particles, at the same time that the particles that are still in the water are being degraded and moved horizontally, and (b) they do not consider time as one of their variables, only depth.

Water bodies are complex systems with varied hydraulic dynamics. Studying the vertical aspects of eDNA in a natural system is a difficult task due to many factors acting in the water column at once (Jane et al. 2015). Flow, hyporheic exchanges, streambeds, surface-subsurface exchange, sediment and colloidal interactions are some of these factors that contribute to this complexity (Shogren et al. 2016; Shogren et al. 2019). Controlling these variables in a field experiment to understand how they affect the vertical dynamics of eDNA is not logistically viable, so they must be studied individually in a controlled environment.

In this study, we aim to understand how free DNA behaves in a controlled water column. To this end, we built a 5-m high and 20-cm diameter PVC tube, injected DNA at the bottom and monitored how it spread through the water column for 8 hours. Understanding how eDNA behaves in the water column is important to interpret species distribution in a water body and improve sampling strategies. A controlled environment is ideal for this because one can introduce variables as our understanding of these dynamics improves.

## Materials and Methods

### Experimental setup

We build an experimental apparatus to emulate the water column of lentic, freshwater conditions using a 5-m high polyvinyl chloride (PVC) pipe (20 cm in diameter). We placed chromatographic septa at six depths (*i.e.* 0, 1, 2, 3, 4, and 5 m) to allow water sampling by the external side of the cylinder by using a medical sterile 1-mL syringes and thus minimizing the generation of turbulence in the water column inside the cylinder. Prior to each experiment, the entire apparatus was decontaminated in a two-step process. First, we used a dichloroisocyanurate solution (0.06 g L^−1^) to thoroughly wash the pipe. We then rinsed away the chlorine with previously treated DNA-free water. This treatment consisted of decontaminating the water with a 10% sodium hypochlorite solution (0.2 mL L^−1^), followed by chlorine neutralization using a 50% sodium thiosulfate solution (0.1 mL L^−1^). The second step was repeated three times in order to ensure that there was no leftover chlorine in the system, coupled with a colorimetric method to detect chlorine after each washing (Zall et al. 1956).

We generated a test solution of eDNA by amplifying a ~100 bp fragment of the COI gene from a genomic sample of the golden mussel *Limnoperna fortunei* (Mytilidae), which is an organism commonly known for its biofouling impacts on hydraulic systems (Darrigran and Damborenea 2011). Each assay was run in a 25 μL final volume reaction, with concentrations: 100 μM each primer, 0.25 mM dNTP mix, 1 U Platinum Taq DNA Polymerase, 1 X Platinum Taq buffer and 2 mM MgCl_2_; Thermocycling conditions followed: 1 min at 95 °C for initial denaturation, followed by 40 cycles of denaturation at 95 °C for 1 min, annealing at 60 °C for 30 sec. and at 70 °C for 30 sec. To obtain a high DNA concentration for the stock test solution, we carried several independent PCRs and the resulting products were pooled, quantified using Qubit 4 fluorometer, and frozen at −80 °C. Immediately before the beginning of the experiments, the DNA solution thawed at room temperature, and each experiment used a 1 mL aliquot (2000 ng of target DNA).

Each experiment began by filling the entire apparatus with DNA-free water up to 5 m, followed by a 15 min period to allow for the water movement to subside. At this point, 1 mL water samples were then collected from each depth using disposable, DNA-free syringes to serve as negative controls. The experimental solution aliquot was then injected at the base of the pipe (5 m depth), and immediately, 1 mL samples were collected from all depths using disposable syringes. Sampling was then repeated at 30 min, 1, 2, 4 and 8 hours after the injection. The entire experiment was run in triplicate. Water samples were then stored at 2 mL decontaminated microtubes, and frozen at −20 °C until analysis.

### eDNA amplification and quantification

Each water sample from the experimental apparatus was processed for DNA extraction using a Solid Phase Reversible Immobilization (SPRI) protocol (DeAngelis et al. 1995). First, 1 mL of collected sample was incubated with a final concentration of 12.5% weight/volume PEG-8000, 0.7 M NaCl and 0.02 mg/mL carboxylated magnetic beads at room temperature for 10 min to condense DNA and bind it onto the magnetic beads. Samples were then magnetized using a neodymium rare earth permanent magnets (NEB), and the supernatant was carefully removed using a micropipette. Samples were dried at room temperature, eluted into 100 μL of TE Buffer and gently mixed. After unbinding DNA from the magnetic beads, samples were magnetized again and the supernatant containing DNA was removed and stored in individual vials.

After extraction, samples were quantified using rtPCR with a hydrolysis probe (TaqMan) targeting the 100 bp fragment previously amplified COI fragment (Pie et al. 2017). Each assay was run in a 10 μL final volume, with concentrations as follows: 75 μM each primer, 25 μM probe and 1 X QuantiNova Probe PCR Kit (Qiagen). Each sample was run in triplicate, with 3 μL of extract being used in each reaction. Cycling conditions were: 2 min at 95 °C for enzyme activation, followed by 50 cycles of denaturation at 95 °C for 5 s, and combined annealing and extension at 60 °C for 5 sec. Assay was run in RotorGeneQ 5plex+HRM (Qiagen). For quantification, a standard curve was built by running a six-order serial dilution of the stock solution previously quantified using Qubit, also performed in triplicate. Each run was analyzed using RotorGeneQ Series Software (Qiagen), with Quantification analysis. Threshold was calculated with automatic option, with a 0.35 upper bound limit, and quantification was done with slope correct mode.

### Analyses

We used two approaches to assess the vertical distribution of eDNA over time. First, we tested the relationship between depth and concentration using linear regressions for each experimental period and determined the time until this relationship became nonsignificant (i.e. DNA concentrations were homogeneous between depths) as an indication of non-homogeneous distribution of eDNA across the apparatus. Second, we fit cubic smoothing splines to each dataset (degrees of freedom = 4). Given that the final concentrations are unlikely to become precisely equal due to measurement error, we compared the observed data to re-sampled splines in which concentrations and depths were randomly shuffled (N=1000 pseudo-replicates). This procedure allowed for the generation of a visual expectation of the expected variation in concentration estimates given the inherent variability of the environmental setup used in our study. All analyses were carried out using R 4.0.2 (R Core Team 2020).

## Results

The vertical distribution of experimental DNA at different time periods is shown in Figure 1. There was a significant relationship between depths and Ct immediately after the beginning of the experiments (t = 2.99, p = 0.008) and after 30 min (t = 5.36, p = 6.32e-05), but that relationship became non-significant after 1 h (p = 0.48 - 0.98). This difference was accompanied by an increase in the DNA concentration across all depths in a manner consistent with the homogenization of DNA concentration throughout the entire apparatus. These results were consistent with the comparison between the splines fit to the observed data and those obtained from shuffled samples. The only two time periods that were outside the simulated data were immediately after and 30 min after the beginning of the experiments. Interestingly, in the latter, the DNA distribution was midway between the state at t=0 and the complete homogenization found at the end of the experiments, with higher-than-expected concentrations up to 3 m from the origin of the DNA.

**Fig. 1.**
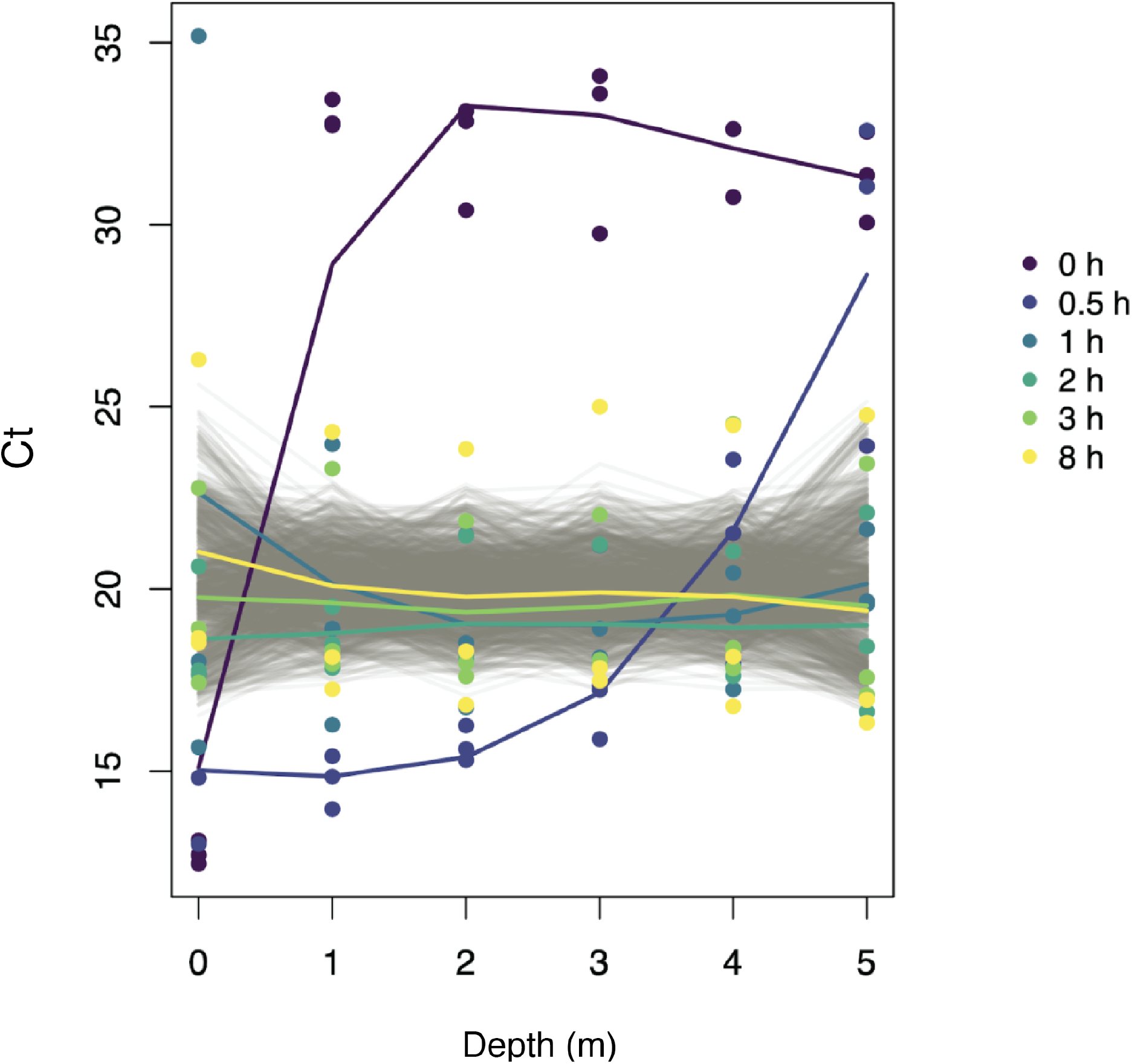
Variation in threshold cycle (Ct) in our experiments from immediately after the addition of DNA (t = 0 h) to eight hours later. Colored lines show cubic splines across the three replicates of each experimental group (see legend). Gray lines indicate 1000 similar splines with datasets in which concentration and depth data were randomly shuffled.

## Discussion

As eDNA studies become increasingly used to monitor different components of the aquatic biota, it is crucial to understand the factors determining the distribution of eDNA in the water column. In our study, we demonstrate that the diffusion of DNA along the water column takes place rapidly, in the time scale of minutes, even in the absence of turbulence. This result is important given that, under field conditions, the water currents would tend to accelerate the homogenization process. Thus, there does not seem to be “an optimal location” to obtain water samples for eDNA analyses in a lentic system, as DNA tends to not accumulate in a specific part of the water column. These results are intriguing, given that previous studies suggested a differential accumulation of eDNA on either the surface (e.g. (Murakami et al. 2019) or the bottom (e.g. (Moyer et al. 2014), or even near the layer that the organism lives (e.g. (Minamoto et al. 2017).

It is important to note that, although we used free DNA molecules in our experiment, eDNA is not a monodisperse phase in nature (Turner et al. 2014; Wilcox et al. 2015). It is composed of particles ranging from single DNA molecules to tissue fragments (e.g. between 0.2 and 180 μm, but mostly between 1-10 μm (Turner et al. 2014)). Particle size composition also plays an important role in how studies comparing different depths report due to how they interact with filter pore size. Although this distribution range seems to be constant between close-related taxa (such as fish, (Barnes et al. 2020)), it seems to vary between different taxa (such as water fleas) (Moushomi et al. 2019). This distribution also changes with time, as bigger particles tend to break down into smaller particles (Murakami et al. 2019). As the sampling and processing methods (volume used, filtration technique, time between sampling and water composition measurements) in these comparative studies are not standardized, it is expected that particle size distribution will play a major role in the results. The behavior of different particle sizes on the water column is unknown. This is a potential source of bias on the sampling, as the captured eDNA can differ significantly from true eDNA source amount on a determined sampling point, because of pore size and volume configuration, and this error can vary between sampling points.

We also expect that the solubility of these different particles influences how they behave in the water column. While most of the eDNA particles tend to have a hydrophilic nature, some are hydrophobic. When considering colloidal particles in the water, eDNA particles can bind to it and behave differently from how they would if they were suspended, mostly due to weight changes. This can lead to accumulation in certain parts of the water column, or changing speed of diffusion (Cai et al. 2006a; Cai et al. 2006b). When bound particles are too dense, it can also promote deposition and accumulation of eDNA in the substrate (Zhai et al. 2019). Size of suspended particles also influences this dynamic, as finer substrates tend to capture more eDNA due to smaller pores (Shogren et al. 2016). This can lead to an effect of accidentally re-suspending trapped eDNA into the water column while sampling, which can cause a sampling bias where capturing water near the bottom is actually capturing the substrate (Turner et al. 2014; Turner 2004). Hydrogeomorphic features of the system being studied should be assessed in order to evaluate slopes (which influence depth variations) and adsorption sites (which can sequester eDNA) (Fremier et al. 2019).

It is also important to emphasize that our results only pertain to a specific aspect of DNA distribution, namely the vertical diffusion process over time in the absence of water currents. The movement of water in lotic conditions might provide qualitatively different conditions, given that water velocity varies with depth. For instance, under laminar flow, water near the surface might include eDNA from farther upstream than those near the bottom (Curtis et al. 2020). However, as water speed becomes faster, the onset of turbulent flow might lead to homogenization of bottom and top water layers (Mächler et al. 2020). Water flow and stratification are also important factors that can create different degrading zones in the water column (Curtis et al. 2020). Liquid flow is known to degrade eDNA due to mechanical forces (Levy et al. 1999). When hyporheic exchanges (water from the main river flow being exchanged with water kept in porous substrates) are considered, we would expect it to create less intense flow zones. These islands could potentially serve as less degrading spaces, where it would be more advantageous to sample near porous substrates both due to sequester of eDNA and due to irreversible sorption to bed sediment (Foppen et al. 2013). Little is known about the dynamic of these spaces regarding eDNA particles and their distribution. In another scenario, when there’s permanent water column stratification (such as in the sea), depth becomes an important sampling factor (Jeunen et al. 2020). It is unknown if eDNA can pass these barriers (i.e. if convection is enough to break these barriers and homogenize eDNA). It’s also unknown if there are clines through the same zones, causing in-between convection to cycle the water and homogenize the water in each water break.

While our results show the behavior of a monodisperse phase of eDNA particles in a relatively small water column, it highlights how this system would behave without interference. With so many factors acting at once in a complex water body system, it is important to break down its components and understand how they behave separately, so we can build a better model that can be incorporated in realistic field conditions. Understanding the interplay between turbulence, colloidal particles and eDNA transport is a particularly important frontier of eDNA research.

## Acknowledgements

This paper presents part of the results of the P&D project, code PD-06491-0383/2015, executed by the Federal University of Paraná and Aliança Prestadora de Serviços Ltda and funded by COPEL Geração e Transmissão SA, under the Research and Technological Development Program of Electricity Sector, regulated by the National Electric Energy Agency (Aneel). The authors declare that they have no conflict of interest.

